# Australia as a global sink for the genetic diversity of avian influenza A virus

**DOI:** 10.1101/2021.11.30.470533

**Authors:** Michelle Wille, Victoria (Tiggy) Grillo, Silvia Ban de Gouvea Pedroso, Graham W. Burgess, Allison Crawley, Celia Dickason, Philip M. Hansbro, Md. Ahasanul Hoque, Paul F Horwood, Peter D Kirkland, Nina Yu-Hsin Kung, Stacey E. Lynch, Sue Martin, Michaela McArthur, Kim O’Riley, Andrew J Read, Simone Warner, Bethany J. Hoye, Simeon Lisovski, Trent Leen, Aeron C. Hurt, Jeff Butler, Ivano Broz, Kelly R. Davies, Patrick Mileto, Matthew Neave, Vicky Stevens, Andrew Breed, Tommy T. Y. Lam, Edward C. Holmes, Marcel Klaassen, Frank Y. K. Wong

## Abstract

Most of our understanding of the ecology and evolution of avian influenza A virus (AIV) in wild birds is derived from studies conducted in the northern hemisphere on waterfowl, with a substantial bias towards dabbling ducks. However, relevant environmental conditions and patterns of avian migration and reproduction are substantially different in the southern hemisphere. Through the sequencing and analysis of 333 unique AIV genomes collected from wild birds collected over 15 years we show that Australia is a global sink for AIV diversity and not integrally linked with the Eurasian gene pool. Rather, AIV are infrequently introduced to Australia, followed by decades of isolated circulation and eventual extinction. The number of co-circulating viral lineages varies per subtype. AIV haemagglutinin (HA) subtypes that are rarely identified at duck-centric study sites (H8-12) had more detected introductions and contemporary co-circulating lineages in Australia. Combined with a lack of duck migration beyond the Australian-Papuan region, these findings suggest introductions by long-distance migratory shorebirds. In addition, we found no evidence of directional or consistent patterns in virus movement across the Australian continent. This feature corresponds to patterns of bird movement, whereby waterfowl have nomadic and erratic rainfall-dependant distributions rather than consistent intra-continental migratory routes. Finally, we detected high levels of virus gene segment reassortment, with a high diversity of AIV genome constellations across years and locations. These data, in addition to those from other studies in Africa and South America, clearly show that patterns of AIV dynamics in the Southern Hemisphere are distinct from those in the temperate north.

**Author Summary:** A result of the ever-growing poultry industry is a dramatic global increase in the incidence of high pathogenicity avian influenza virus outbreaks. In contrast, wild birds are believed to be the main reservoir for low pathogenic avian influenza A virus. Due to intensive research and surveillance of AIV in waterfowl in the Northern Hemisphere, we have a better understanding of AIV ecology and evolution in that region compared to the Southern Hemisphere, which are characterised by different patterns of avian migration and ecological conditions. We analysed 333 unique AIV genomes collected from wild birds in Australia to understand how Australia fits into global AIV dynamics and how viruses are maintained and dispersed within the continent of Australia. We show that the Southern Hemisphere experiences differing evolutionary dynamics to those seen in Northern Hemisphere with Australia representing a global sink for AIV.

## Introduction

The evolution of avian influenza virus (AIV) is in part driven by the globally booming poultry industry that comprises an estimated three quarters of the global avian biomass [1, 2]. This industry has witnessed a dramatic increase in the incidence of disease outbreaks over the past two decades caused by high pathogenicity avian influenza virus (HPAIV) [3, 4]. Despite this, wild birds continue to play an important role in AIV ecology and evolution. Through long distance migration, wild birds have aided in the dispersal of high pathogenicity H5Nx between Asia, Europe, Africa and North America [5]. Conversely, the existence of distinct migratory flyways has constrained viruses into consistent phylogenetic divisions, such as between AIVs detected in the Nearctic and the Palearctic [6]. Aggregations of wild birds vary both geospatially and temporally, often leaving their hallmarks on AIV prevalence, diversity and evolution [7]. Importantly, most of our knowledge of AIV ecology and evolution is drawn from studies in temperate northern hemisphere systems [8–15] even though migration patterns and environmental conditions relevant for AIV dynamics differ in the southern hemisphere [7].

Influenza A virus is a segmented, negative-sense RNA virus and the sole member of the genus *Alphainfluenza* in the family *Orthomyxoviridae* [16]. Wild birds, particularly *Anseriformes* (ducks, geese and swans), and to a lesser extent *Charadriiformes* (shorebirds and gulls), are central reservoirs of AIV, with 16 of the 18 HA (haemagglutinin) and 9 of the 11 NA (neuraminidase) subtypes identified in these taxa [6, 17, 18]. AIVs do not generally cause high morbidity or mortality in their hosts, with the exception of subtype H5 and H7 HPAIVs that emerge in poultry [5, 19–21]. In northern hemisphere systems, the prevalence of AIV peaks in the autumn, driven by the recruitment of immunologically naïve juvenile avian hosts and population congregations associated with migration [8, 9, 18, 22]. However, disease dynamics may vary in different global regions due to differences in environmental and host factors [23]. Indeed, despite many parts of Australia being defined as temperate, annual recruitment of immunologically naïve juvenile waterfowl into avian populations is irregular due to highly variable climatic conditions which impact breeding cycles, such that in some years waterfowl may not breed, or breed in small numbers and in other years may attempt to breed multiple times [24]. Unlike the high prevalence of AIVs in temperate northern hemisphere waterfowl, prevalence in Australian waterfowl has consistently been less than 2% with no strong seasonal patterns, however these low prevalence estimates may be driven by the highly aggregated nature of studies [25–32]. Furthermore, all Australian waterfowl are endemic and largely nomadic, and do not migrate beyond the Australian-Papuan Region [33, 34]. Indeed, of the key AIV reservoir avian taxa, only members of the *Charadriiformes*, notably the waders (families *Scolopacidae* and *Charadriidae*), migrate and link Australia with Eurasia and North America [35–37]. These species may also be less susceptible to AIV infection than some other species [38]. The ecology of this migratory system has the potential to limit viral gene flow between Eurasia and Australia and, consequently, it is expected that AIV lineages may be evolving independently in Australia compared to other continents [27, 39]. Aside from a small number of studies based on a limited number of AIV sequences [27, 30, 31, 39–43], how these distinct features of host-ecology impact AIV evolution in Australia is largely unknown.

To reveal patterns of AIV evolution in wild birds in Australia, we used low pathogenic AIV (LPAIV) genome data collected over nearly 15 years from all states and territories of Australia to assess (i) the pattern of gene flow between Australia and other continents i.e. a source, sink or combination, (ii) the extent and role played by AIV lineage introduction and maintenance in Australia *e.g.* local evolution and extinction, and (iii) whether the population dynamics, migration and reassortment of AIVs in Australia differ from those in other geographical locations globally. Studying these processes in Australia, which comprises conditions that are in stark contrast to those found in temperate avian population systems in the northern hemisphere, will provide key insights into the global drivers of AIV ecology and evolution.

## Results

### Summary of avian influenza viruses sequenced

The data generated here comprised full or partial genomes of 333 unique LPAIV. Briefly, a total of 397 AIV positive samples collected from 2006 to 2020 were submitted for sequencing (Table S1). We recovered the full AIV genomes from 242 of the samples. In some cases, we recovered partial AIV genomes consisting of gene segments with insufficient sequence length (n=15) or with no sequence (n=76). An analysis of the influence of Ct value and genome completeness is presented in Fig S1. A small number of the virus samples comprised mixed infections (n=20), where two different variants of a segment were detected. Forty-five viruses were sequenced more than once; samples may have been re-sequenced due to poor quality in the initial attempt and/or in cases where both the original sample and the corresponding egg isolate were sequenced.. Our analysis also included additional virus sequences presented in Bhatta *et al.* 2020 (n = 1, individual avian faecal sample in 2018) and Hoye *et al.* 2021(n = 22, combined oropharyngeal cloacal swabs collected in 2014) as the samples were collected as part of the NAIWB surveillance program.

Overall, unique AIV genomes comprising at least one segment characterised were collected in South Australia (n = 71), Western Australia (n = 75), Tasmania (n = 89, including [41], Queensland (n = 45), Victoria (n = 46, including [40], New South Wales (n = 15) and the Northern Territory (n = 15) (Fig 1A). These include those collected from avian cloacal and/or oropharyngeal swab samples or avian faecal samples, and unique genomes were generated from a combination of original samples (n=222) or isolates (n=111). Prior to 2013, there were fewer than five sequenced genomes per year. However, since this time the numbers of virus genomes have steadily increased, with the largest number of genomes sequenced from samples collected in 2019 (n = 71) (Fig 1). This increase coincided with a shift by the NAIWB surveillance program from characterising only H5/H7 viruses towards more comprehensive LPAIV characterisation in Australia. Due to irregular data collection in some states, large numbers of viral genomes were recovered from single sampling events (*e.g.* Western Australia, Tasmania), whereas in other states we find a more uniform temporal spread of the data (*e.g.* Victoria) (Fig S2).

**Figure 1.**
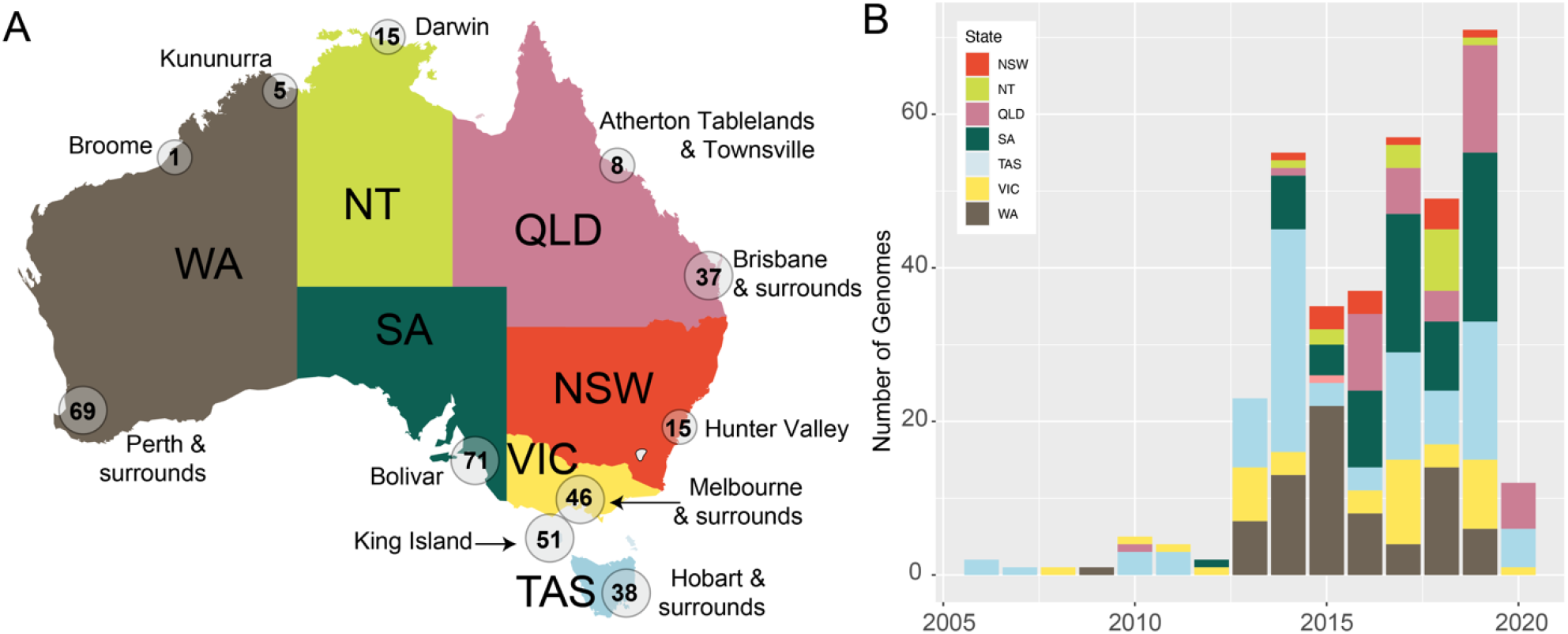
Spatial and temporal distribution of avian influenza genomes used in this study. (A) Map of Australia illustrating regional sampling locations. Where sampling locations were within 500km, they were merged into a single location. The value within the circle corresponds to the number of unique viral genomes comprising at least one segment from each location. States and Territories are as follows: VIC Victoria, NSW New South Wales, QLD Queensland, NT Northern Territory, WA Western Australia, SA South Australia and TAS Tasmania. (B) Number of genomes per state per year. Colours from panel B correspond to the fill colour of the state in panel A. This figure includes genomes comprising one or more segments and contains no duplicates. This figure includes all sequences generated as part of the National Avian Influenza Wild Bird Surveillance Program, including those recently published in [40, 41]. Metadata is available in Table S1 and a detailed plot illustrating exact virus sample collection dates and locations can be found in Fig S2.

Across the data set as a whole, we identified 14 different HA subtypes and all nine different NA subtypes, comprising 58 HA-NA combinations. We did not detect avian HA subtypes H14 and H15, and only a single case each of H13 and H16. The most common subtypes in our data set were H1N1 (n = 14), H3N8 (n = 23), H4N6 (n = 16), H5N3 (n = 15), H6N2 (n = 23), H9N2 (n = 14) and H11N9 (n =19) (Fig 2A). These subtypes each comprised 5-10% of the subtype combinations. An analysis of HA-NA linkage by assessing the Pearson’s residuals following a Chi-squared test revealed a strong positive association between H1-N1, H3-N8, H4-N6, H8-N4, H11-N9 and H12-N5 (Fig 2B). These overrepresented subtypes are comparable to those recovered from intensively sampled study sites in Europe and North America [8, 10]. In cases in which a HA subtype had several different NA subtypes, we saw weak positive or weak negative Pearson residuals (*e.g.* H7). As our data set largely comprised samples collected from wild bird faeces and pooled samples, the contribution of avian host species to AIV subtype distribution could not be determined.

**Figure 2.**
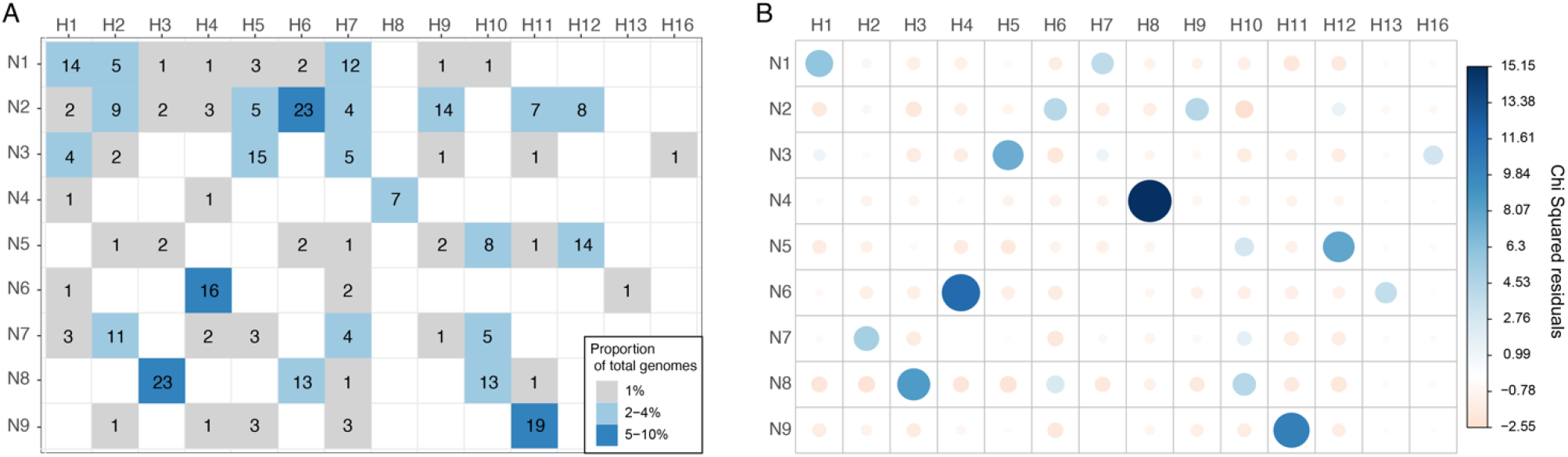
HA-NA subtype linkage in data generated for this study. (A) The number of each HA-NA subtype combinations (values) and the proportion of the total data set these values represent (shading). (B) A plot of the Pearson residuals of Chi-squared tests. For a given cell, the size of the circle is proportional to the amount of the cell contribution. Positive residuals are in blue and identify HA and NA subtypes for which there is a strong positive association in the data set. Negative residuals are in light pink and show a weak negative association, that is, they are underrepresented in the data set. This figure comprises unique viral genomes with at least one segment.

### Australia is a sink for global AIV diversity

Using the data generated here, we first aimed to determine how AIVs in Australia fit into patterns of global genetic diversity. Specifically, we asked (i) whether there was one consistent endemic Australian lineage for each subtype, (ii) how long these endemic lineages have been maintained, and (ii) whether there is connectivity between Australia and New Zealand (sequences mined from GenBank) comprising an “Oceania cluster” within the southern hemisphere temperate zone.

Our phylogenetic analysis revealed that sequences from Australia tended to fall into distinct Australian lineages, although the number of these lineages varied across subtypes and segments. In the case of the HA segment, sequences from H2-H8 subtypes each comprised a single contemporary lineage (Fig 3, Fig S4-S9). In contrast, H1, H9-H12 had more than one contemporary lineage (Fig 3, Fig S3, S10-S12); Subtypes H9 had three, and H1, H10-H12 each had two, contemporary lineages. In addition to lineages, there was evidence of at least one H10 and two H11 incursions into Australia without subsequent establishment (Fig 3, Fig S11). These differences in the number of lineages concur with the observation that H1-H6 are over-represented at duck-focused study sites as compared to H8-H12 which are under-represented at duck-focused study sites [*e.g.* 8]. It has been proposed that ducks may be not be the central reservoir for H8-H12 [44]; based on phylogenies (Fig 3, Fig S9-S12) there was evidence for more repeated incursions of these subtypes into Australia.

**Figure 3.**
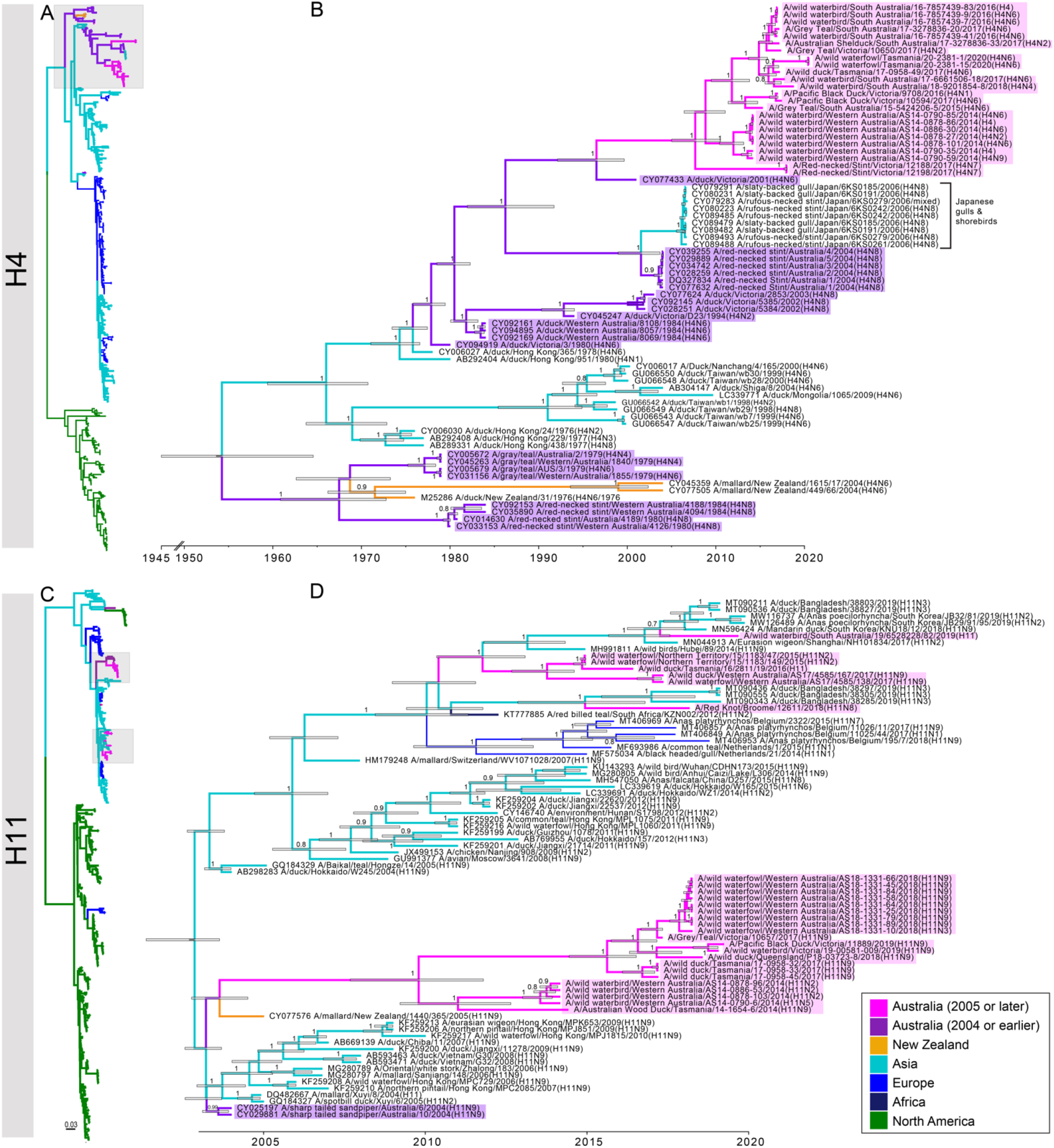
Phylogenetic trees of subtypes H4 and H11. (A, C) Phylogenetic trees comprising global diversity. Branches of reference sequences are coloured by continent. Sequences from Australia are coloured in pink (2005 and later) and in purple (pre-2005), with 2005 marking the year of the oldest sequence in the data set generated in this study. (B, D) Time structured phylogenetic trees. The trees comprise Australian lineages (as indicated by grey boxes in A, C) and closest relatives (retrieved by BLAST searches). Branches are coloured by year and geography as above. Branch labels correspond to posterior clade probabilities of each node, node bars correspond to the 95% HPD of node height. We selected H4 as it is the only HA subtype for which there is clear introduction of an Australian lineage virus into Asia (indicated in square parenthesis), and is an example of an HA segment for which there is only a single contemporary lineage. We selected H11 as it is the subtype with the largest number of contemporary Australian lineages (4), of which 2 are represented by a single sequence. Trees for all other HA subtypes can be found in Fig S3-S14

For many lineages in the time-scaled trees, there was a relatively large time gap between the most recent common ancestor of Australian lineages and the closest reference sequence (Fig 4). For example, in the Australian H1 lineage represented by four viruses from 2013 and 2016 (Fig S3), the time to the most recent common ancestor (tMRCA) ranged from Feb 2011-June 2013 (95% Highest Posterior Density [HPD]; mean at June 2012), whereas their date of separation from the closest reference sequence was between 1999-2003 (mean at June 2001) (Fig S3). This is most likely due to vast under-sampling in Australia, notably between 2000 and 2012, although sporadic and/or under-sampling of wild birds in Asia may compound this. Critically, this has implications for accurate dating of some Australian lineages as it is unclear how distant the introduction of the lineage to Australia predated the tMRCA of existing diversity (Fig 4). These issues notwithstanding, the tMRCA of contemporary Australian lineages was 2005 or later, suggesting currently established lineages were introduced to Australia relatively recently (Fig 4). This was supported by the fact that most of the older Australian lineages, comprising viruses from the 1970s to 1980s are no longer in circulation. There are some exceptions, such as H7 viruses, that had a tMRCA of between Aug 1974 - Aug 1975. This Australian H7 lineage has been associated with eight HPAIV poultry outbreaks in Australia since 1976 [45]. Sequence data for this H7 lineage from wild birds has only been available since 2007 due to very limited sampling and sequencing of wild birds in earlier years (Fig S8).

**Figure 4.**
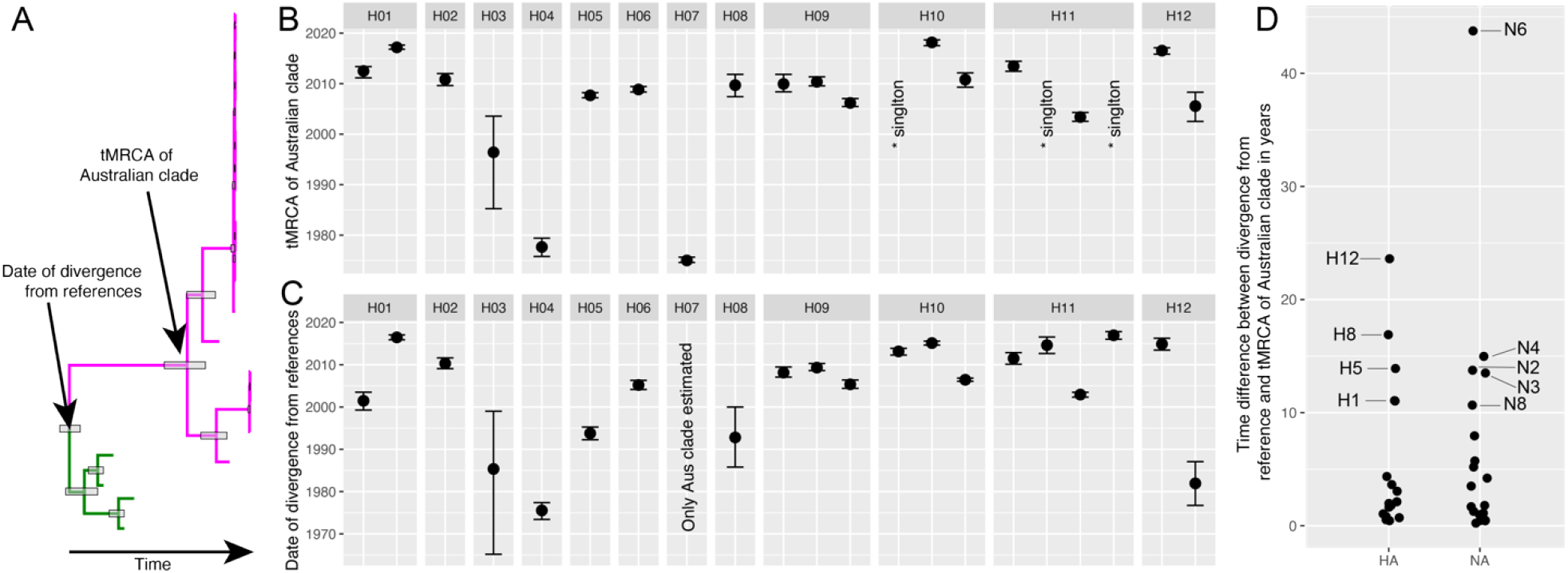
The time-scale of AIV evolution by subtype in Australia. (A) A schematic phylogeny demonstrating the differences between the tMRCA of Australian lineages and the dates of divergence from reference sequences. (B) The tMRCA distribution of contemporary Australian lineages of the HA segments and (C) dates of divergence from the reference sequences of all HA lineages. Points represent the node date and bars the 95% HPD. For segments with multiple lineages, multiple estimates have been provided. Where a novel introduction is represented by a single sequence the tMRCA was not estimated (here, represented by “*singleton) but the date of divergence from reference sequences is shown. For H7, we did not estimate the date of divergence from the closest reference sequences. (D) The time difference between the tMRCA of the Australian lineages and the date of separation for all HA and NA segments and lineages. Lineages with time differences of more than 10 years are labelled. All HA and NA trees are presented in Fig 3, Fig S3-S23.

Notably, sequences from Australia and New Zealand did not consistently fall into the same lineages. There were 42 HA sequences from New Zealand available in GenBank, of which 33 had collection dates of 2004 and later, aligning well with the temporal scale of contemporary Australia lineages. Indeed, New Zealand lineages of H1, H2, H5, H6, H7 and H10 were each in entirely separate lineages from Australian sequences (Fig S3-S4, S6-S8, S11). This is likely due to limited bird migration involving some shorebird species, between New Zealand and Australia [46].

Overall, Australia appears to be a sink for Eurasian AIV diversity. Although we identified multiple viral introductions from Eurasia to Australia, in the entire data set there were only two examples of viruses from Australia being introduced to Eurasia. These comprised the H4(N8) subtype (Fig 3B), and one N7 sequence (Fig S21). Notably, each of these events involved the detection of Australian lineage viruses in Charadriiformes (gulls and shorebirds) in Japan. Overall, all viral introductions stemmed from Eurasian lineages with the exception of H10 and H12 that showed introductions from North American lineages. For H8 and one H9 lineage, the most closely related reference viruses were sampled in Europe. However, due to possible under-sampling and/or under-representation of viral diversity in wild birds in Asia it cannot be concluded that these lineages were seeded directly from Europe (Fig S9-S10).

### Detection of novel virus segments introduced into Australia

The relatively long time difference between the tMRCA of Australian lineages and the global representative viruses used as reference suggest that the wild bird surveillance sampling has been unable to detect the index viruses seeding local lineages (Fig 4). However, a small number of viruses in the data set (n = 18) contained gene lineages and/ or introductions with no further transmission which likely comprise recent introductions to Australia. These viruses comprised at least one virus gene segment that either represented the only detection of a novel lineage in Australia (*i.e.* singletons) or comprised the first detection of an Australian lineage cluster, where the time difference between the tMRCA of the identified lineage and date of divergence from global references was small (less than 1 year) (Fig 5). These recently introduced viruses were only detected in the north of Western Australia, the Northern Territory and Queensland, and from migratory shorebirds in Tasmania. Migratory birds would likely use these northern locations as initial stopover sites in Australia, highlighting the importance of surveillance of shorebirds in these regions. Notably, we did not find evidence of a complete “novel” virus genome, that is all viruses for which whole genome data were available contained at least one gene segment belonging to an established Australian lineage. For example, A/Ruddy Turnstone/King Island/10938/2017(H12N5) had 7 segments representing the index detection of a novel lineage, with only the M segment belonging to an established Australian lineage. Interestingly, Ruddy Turnstone viruses in 2018 and 2019 had a number of segments falling into lineages for which A/Ruddy Turnstone/King Island/10938/2017(H12N5) was basal.

**Figure 5.**
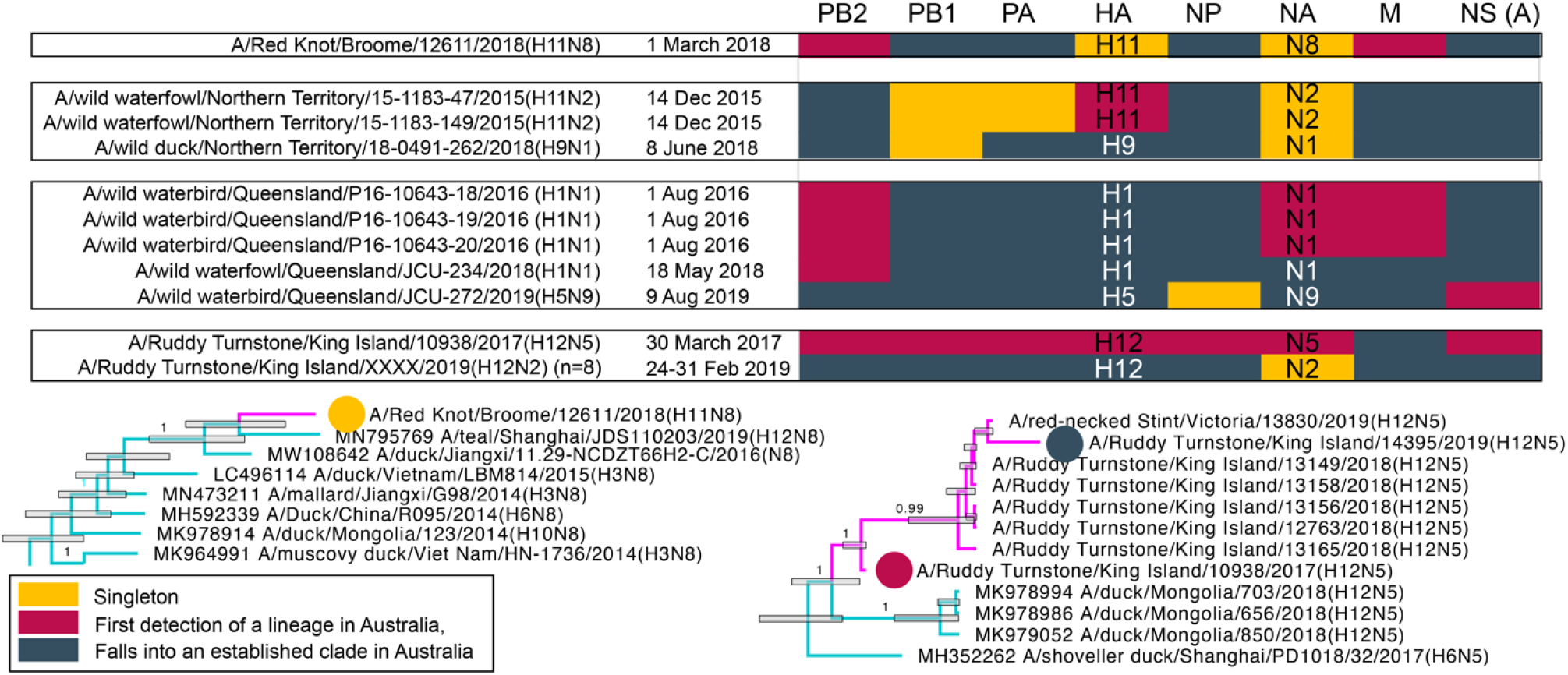
Viruses sequenced in this study that have signatures of recent introduction. For each segment, coloured tiles correspond to three different statuses: singletons, first detections and well-established lineages. Singletons represent the only detections of the lineage in Australia. In cases where two viruses from the same sampling effort were identical and were the only detections of that lineage, they were still considered a singleton (e.g. the NA segment of the NT/2015 viruses or the NA segment of the 8 H12N2 Ruddy Turnstone viruses from 2019, which have only ever been detected during that sampling event). Only the A allele NS segment was detected in these viruses. Phylogenetic examples (here excerpts from N8 and N5) are provided for each status, and branches are coloured as in Fig 3.

### AIV circulation within Australia

Given that there are no structured flyways within Australia and birds have nomadic movements influenced by climate [29, 47], we hypothesized that there would be limited geographic structure of AIVs within the continent. To assess this, we analysed possible viral “migration events” (Markov Jumps) between sampled locations using the HA and/or NA segments comprising Australian lineages with 20 or more sequences (H4, H5, H6, H7, N6, N8), and two independent lineages of the NP segment. These analyses are likely strongly influenced by both small sample sizes and collection biases, such that we do not have sequence coverage across all subtypes for all locations and years. However, we also examined the two larger NP lineages which included substantially more sequences (n = 197 and n = 85 sequences) than any of the subtype specific HA or NA data sets and spanned the entire sampling period and all sample locations.

Our phylogenetic data revealed potential virus migration events between the sampling locations in the southeastern states (Victoria, South Australia, New South Wales and Tasmania) that occurred consistently across all of the gene segments examined (Fig 6). Using the largest Australian NP gene lineage, we found more than 10 potential migration events between Victoria and South Australia and between Victoria and Tasmania, suggesting high levels of connectivity between these sampling locations. We also found evidence of movement between temperate Western Australia and the southeastern states (Victoria, South Australia, Tasmania), and between Queensland and the southeastern states, although this was only detected in the NP segments and in two of the HA/NA subtypes analysed. As only limited sequences were available from tropical Australia (northern Queensland, Northern Territory and northeastern Western Australia), migration events to/from these locations were not well estimated in our analyses. However, for the largest NP lineage, a number of potential migration events between temperate and tropical Australia were observed (Fig S26). Potential migration events were also detected between the sampled tropical locations. Although it is likely that we have underestimated the migration events due to poor temporal and spatial coverage, the migration events had strong Bayes Factor support (Fig S25). Importantly, these analyses also did not record >1 migration event or >10 Bayes Factor between all locations that were included in each tree as a default. For example, despite being included in all eight analyses, we only detected significant migration events (or >10 Bayes Factor) to/from Western Australia in the H6 and N6 lineages, and the two NP lineages (Fig 6, Fig S25).

**Figure 6.**
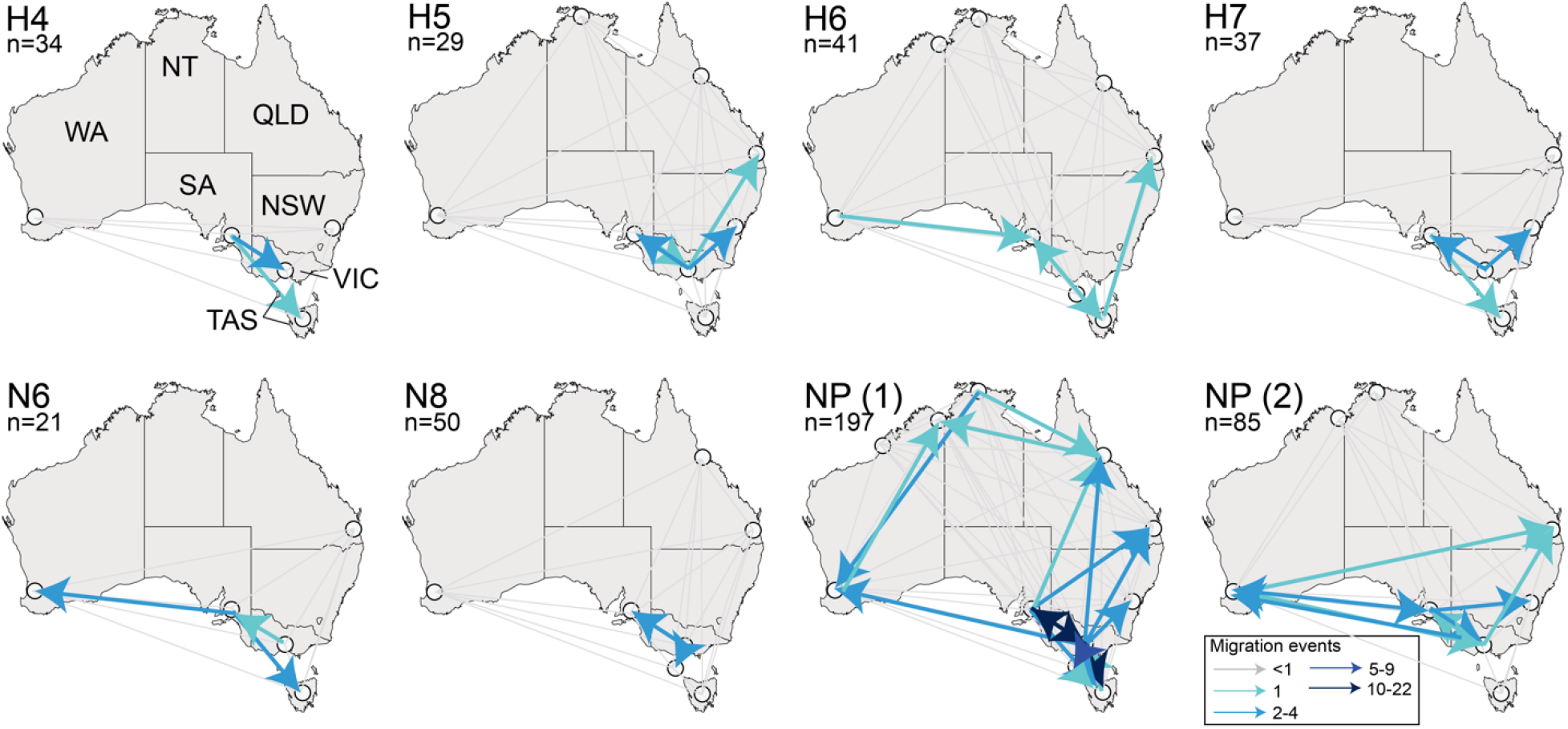
Inferred migration events of avian influenza viruses within Australia. Locations included in each tree are marked by a white circle. Specific location names are presented in Figure 1A, and all state names are presented in the first panel and are as follows: VIC Victoria, NSW New South Wales, QLD Queensland, NT Northern Territory, WA Western Australia, SA South Australia and TAS Tasmania. Grey lines correspond potential migration events that were not detected in the analysis (i.e. migration event <1). Blue lines indicate migration events are derived from calculations of state changes (Markov Jumps), ranging from light to dark. Arrows indicate the direction of the migration event. As NP has more than a single discrete Australian lineage, we have generated two independent maps reflecting the 2 largest Australian lineages of NP (Fig S25). Maps illustrating Bayes Factors, also generated using BSSVS can be found in Fig S26, and Markov rewards also generated in this analysis are presented in Fig S27.

Overall, analysis suggested Victoria was consistently a net exporter as most migration events originated from the state. Specifically, Victoria played a role as a net exporter in H5, H7, N6, N8, and both NP lineages. South Australia also played a role as an exporter (H4, H6, H7, N6, and both NP lineages), although we detected both import and export events from this state across most analyses. Temperate Western Australia was a net importer of AIV, although as with South Australia, we detected both importation and exportation events across the analyses. A positive association between Markov rewards and the number of exportation events may also be evidence of sampling bias. For example, in the case of H4, H5 and H7, Victoria had substantially more sequences available as compared to other sampling locations and was identified as a net exporter. In these cases, the high number of exportations relative to importation events may be due to sampling biases (Fig S27).

Taking the potential biases in our data set into consideration we did not see consistent source or sink locations for AIV movements but rather detected numerous exportation and importation events in most locations. Further, there was no consistent directionality to patterns of viral dispersal. Rather adjacent locations from which we had many samples were highly connected. These results are consistent with the absence of flyway structure within Australia.

### Genomic Reassortment

Despite a low reported prevalence, multiple lineages and subtypes co-circulated at most of the sampled locations (Fig S28), [*e.g.* 25]. A number of “mixed” virus samples (*i.e.* samples comprising at least one segment with two different sequences) were also detected through sequencing. These mixed virus samples were often detected from sampling events where a diversity of AIV subtypes were co-detected. The only exceptions were A/wild duck/New South Wales/M15-10737-MD02/2015(mixed) and A/wild waterfowl/Queensland/JCU-78-226/2016(mixed) for which other AIV genomes were not detected in birds collected on the same collection events (Table S2). Next-generation sequencing of the original samples allows for the detection of mixed viruses.

Assessment of the diversity of genome constellations indicated prolific reassortment, similar to that found in other locations that have been studied [13, 14]. In the case of the H5 and H7 subtypes of veterinary importance, the LPAIV genome data from wild birds revealed 17 unique constellations from 33 (26 complete) H7 genomes, and 18 unique constellations from 29 (20 complete) H5 genomes (Fig 7). The only virus samples with identical genome constellations were those from the same sampling event and location. However, even within the same sampling event where the same HA-NA subtype combination was detected, there was evidence of genetic reassortment. For example, of the 11 H7N1 virus samples sequenced from a single 2019 sampling event in South Australia, six viruses had an NS B allele while the others had the NS A allele (Fig 7). We found that within the same year, partial genome constellations were shared. For example, in 2018 H7 viruses were collected in New South Wales, Queensland, South Australia, and Victoria. With the exception of A/wild waterbird/South Australia/18-7728954-65/2018(H7N6) these viruses share 5 of 8 segment lineages, with differences in PA, NA and NS. (Fig 7).

**Figure 7.**
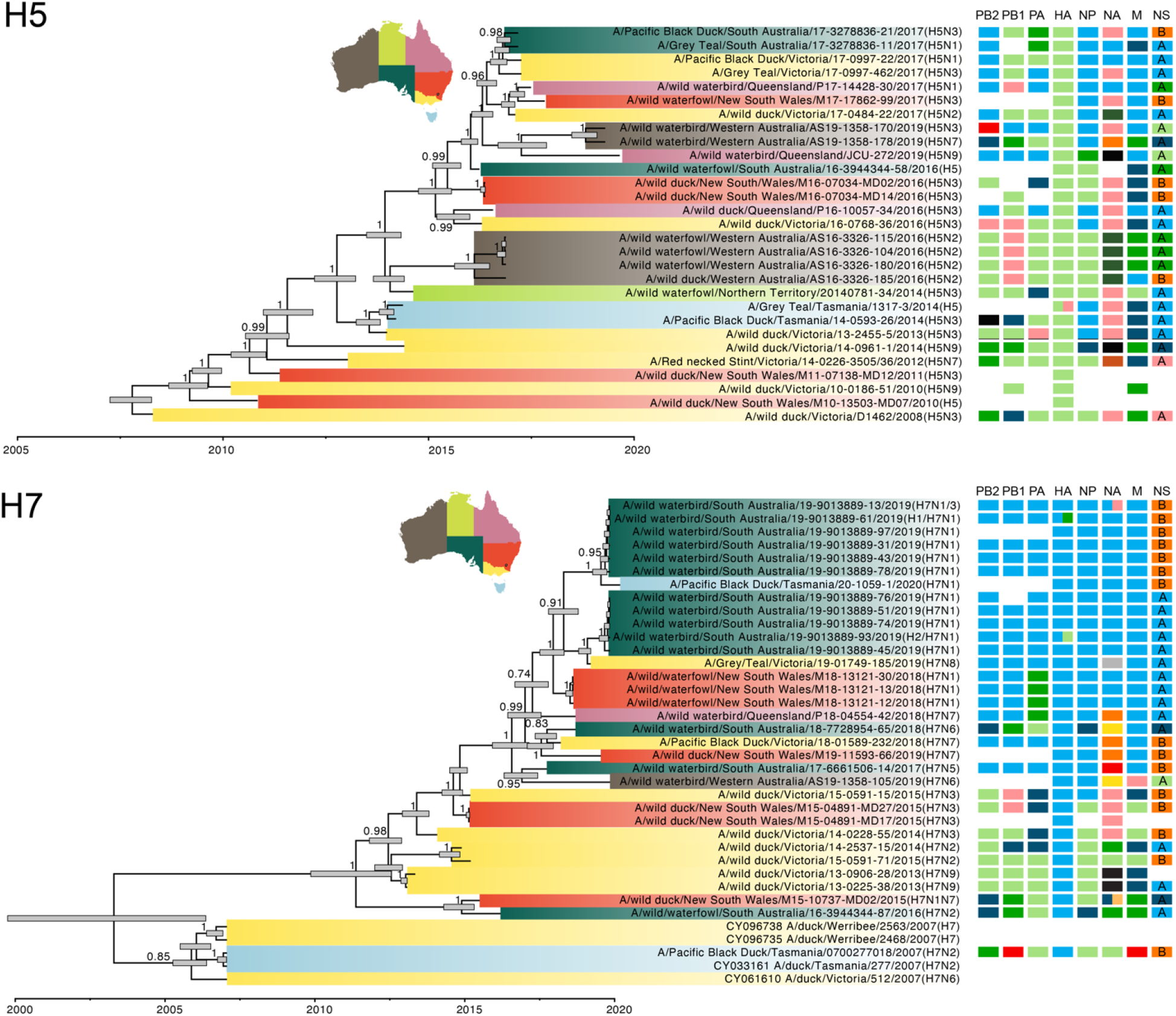
Genome constellations of (A) H5 and (B) H7 viruses. The phylogenies presented are time-scaled Maximum Clade Credibility Trees. Tips are coloured according to Australian state or territory. Scale bar denotes the year of sample collection. Node bars are the 95% HPD of node height, and posterior clade probability is presented on each branch. Adjacent to each tree are coloured tiles where each column of tiles refers to a segment, arranged according to size: PB2, PB1, PA, HA, NP, M, NS. We only included tiles for viruses sequenced in this study and in cases where the tiles are blank, no sequence was available for the segment. Different colours refer to different lineages, whereby tile colour scheme is retained for both H5 and H7 trees. For example, for the NS segment, viruses with an NS B lineage are coloured in orange. The viruses here fall into five different lineage clusters of NS A, and these are presented in two different shades of blue and green and pink. If a virus is a “mixed” infection, segments with two different lineages or subtypes are split to illustrate this.

## Discussion

Australia and the Southern Hemisphere have a chequered legacy for research on avian influenza. The first detection of AIV in wild birds was associated with a mortality event in terns off the coast of South Africa [48], and the first descriptions of avian influenza from wild, healthy birds was from the Great Barrier Reef islands of Australia [49–51]. Furthermore, one of the most enigmatic subtypes, H15, was initially described in Australia [52, 53]. Through surveillance activities, particularly since 2006, it has become clear that, unlike the northern hemisphere, AIV prevalence in Australia is generally low with no strong seasonal pattern, however prevalence estimates generated from current surveillance methods have large uncertainty. Despite low isolation success, recent studies have demonstrated that AIV detections fluctuated temporally and geographically, and that the full diversity of AIV subtypes circulate on the continent [25, 27, 28, 32, 54]. Early phylogenetic studies on a limited selection of AIV subtypes and sporadic sequence data suggested the potential for Australian specific lineages, and detections of intercontinental reassortants [30, 39, 41–43]. Despite these findings, understanding of AIV evolution in wild bird populations of the southern hemisphere lags behind that of the northern hemisphere due to low sampling rates and characterisation of virus data [54, 55]. This study is the first to comprehensively assess AIV evolution across all detected subtypes in Australia, the outcomes of which demonstrate the importance of the globally varying characteristics of bird migration on AIV dynamics.

A key observation was that AIV in Australia are characterised by infrequent enduring introductions followed by decades of isolated circulation. Hence, Australia appears to be a sink for AIV genetic diversity and not closely linked to the Eurasian virus gene pool. This dynamic is mirrored in Africa and South America. Although southern hemisphere AIV lineages sit within lineages originating in the northern hemisphere, our results reinforce findings from a growing number of studies demonstrating that AIV lineages from the temperate north are sporadically introduced to the southern hemisphere [41–43, 56–59]. Specifically, sequences generated in the Neotropics fall into lineages within the Nearctic lineage, and Afrotropical and Australasian lineages are generally part of the Palearctic lineage. In contrast, export events from the southern hemisphere into the temperate north have very rarely been reported [57, 60]. Once introduced, lineages circulate in isolation in the southern hemisphere until extinction. Data from both South America and Australia illustrate that in some cases, lineages have been maintained in isolation for decades. Rimondi *et al* 2018 reported that a unique PB2 lineage has been circulating in South America for ~ 100 years. Similarly, lineages such as those of the Australian H7 subtype viruses, have been circulating in the country continent for more than 50 years, although precisely dating the divergence of these lineages is challenging due to the sparsity of AIV sequence data prior to 1980. Despite long-term isolation, we demonstrated that many lineages that circulated in the 1980’s have become extinct and have been replaced, perhaps due to competitive exclusion as seen in other locations [61].

Waterfowl migration influences viral evolution, and Australia as a sink for AIV diversity is likely driven by a lack of waterfowl migration between Australia and Asia, particularly across the Wallace Line [33]. AIV are predominately distributed by waterfowl, with key evidence described for the flyway system of North America [62], the rapid movement of HPAIV H5Nx viruses across the globe coinciding with waterfowl migration patterns [5], and prevalence peaks in Africa coinciding with the arrival of migratory Palearctic waterfowl [63]. In Oceania, waterfowl species are endemic to the Australo-Papuan region [33]. It is therefore more likely that the limited introduction of novel AIV lineages to Australia are due to long-distance migratory shorebirds flying from their northern hemisphere breeding grounds along the East Asian Australasian Flyway to Australia for the duration of their non-breeding season [64]. This is notably reflected in the larger number of novel introductions detected in subtypes such as H9, H10 and H11, and fewer detectable introductions of subtypes typically associated with waterfowl, especially ducks (*e.g.* H4). Our finding of virus gene lineages originating from both Eurasia and North America further supports that migratory shorebirds play a key role in introducing AIVs into Australia. Alaska is part of the East Asian Australasian flyway [64] and shorebird species such as Sharp-tailed Sandpipers (*Calidris acuminata*) [65], Ruddy Turnstone (*Arenaria interpres*) [35] and Bar-tailed Godwit (*Limosa lapponica*) [66] migrate from Alaska to Oceania. AIV genomes that we identified to contain novel viral introductions were detected in shorebird samples including from locations where shorebirds may stop during southward migration, such as Broome, Western Australia [67]. Like Australia, the Amazon rainforest forms a major barrier to waterfowl migration in the Nearctic-Neotropical system [68, 69]. As such, in a similar manner to Australia, the movement of AIVs from the Nearctic to the Neotropics is mostly likely carried by long-distance shorebird migrants [57, 70, 71]. A large evolutionary study similarly showed the importance of shorebirds in introducing viruses to South America, as many of the recently characterized virus detections from shorebirds belonged to the main North American shorebird-associated lineages rather than divergent South American lineages [57].

Within Australia, we found no evidence of directionality in the movement of the AIV gene pool within Australia. As Australian waterfowl are nomadic rather than strictly migratory, there are no key migratory flyways within the continent. Rather, ducks have “erratic” movement patterns across the continent which are heavily dictated by the availability of water [47]. Therefore, consistent patterns of AIV movement between specific locations would not be expected. However, we did observe high connectivity (*i.e.* the number of strongly supported viral migration events detected within phylogenies) between the southeastern locations. Across this region, movements of waterbirds tracking water within and between the large Murray-Darling and Lake-Eyre basins may form the natural links. For example, satellite tagged Grey Teals (*Anas gracilis*) moved widely across the Murray-Darling basin, utilizing permanent and temporary watercourses in Victoria, New South Wales, and South Australia. Some of these tagged individuals connected the Murray-Darling with sites in Queensland and Northern Territory with flights of over 1200km [47]. While ducks are likely the major driver of virus movement within Australia, there are also a number of nomadic waders that similarly move long distances in search of water for breeding and foraging [72]. Unfortunately, due to low prevalence and a sampling regime not designed to investigate these dynamics, we were unable to infer the fine scale patterns of virus movement.

Our data and analyses are central for placing future Australian AIV genome sequences and studies within the local and global context. Some recent studies have reported the possible detection of novel intercontinental reassortants where AIV segments were reported to be more closely related to lineages from Eurasia and North America compared to those from Australia [40, 41]. Here we clarified that these viruses are not necessarily recent intercontinental reassortments but belong to pre-established lineages in Australia [40, 41]. For appropriate outbreak response and biosecurity policy development it is crucial to accurately assign the source of AIV detected in poultry or wild birds as potential novel introductions of “exotic” viruses or their derivative reassortants, especially in the presence of reassortment promiscuous lineages such as clade 2.3.4.4 HPAIV H5Nx associated with current epizootics in the Northern Hemisphere that may have devastating consequences for the local poultry industry. This, combined with the potential roles of shorebirds in introducing AIV lineages [37] to Australia has implications for wild bird AIV surveillance and risk assessments for wild bird, poultry and human health.

In sum, we revealed that the evolution of AIV in Australia differs from patterns found in the northern hemisphere. These reflect differences in environmental conditions influencing bird ecology, notably in AIV host competency and movement patterns, and taken together should be integrated into improved risk assessments of potential AIV spillover into poultry and the distribution of exotic or potentially zoonotic AIV lineages into Australia.

## Methods

### Ethics Statement

All capture and sampling of wild birds carried out by Deakin University was conducted under approval of Deakin University Animal Ethics Committee (permit numbers A113-2010, B37-2013, B43-2016, B39-2019, B03-2020), Philip Island Nature Park Animal Ethics Committee (SPFL20082), Wildlife Ethics Committee of South Australia (2011/1, 2012/35,2013/11) and Department of Primary Industries, Parks, Water & Environment Animal Ethics Committee of the Tasmanian Government (5/2019-20). Banding was done under Australian Bird Banding Scheme permit (banding authority numbers 2915, 8000, 8001). Research permits were approved by Department of Environment, Land, Water and Planning Victoria (10005726, 10006663, 10007534, 10008206, 10009534), Department of Primary Industries, Parks, Water & Environment of the Tasmanian Government (FA11255, FA13032, FA14110, FA15270, TFA14065, TFA15269, TFA16256, TFA17018, TFA18088, TFA19044), Office of Environment and Heritage New-South Wales (SL101252), Department of Environment, Water and Natural Resources South Australia (M25919-1,2,3,4,5), Department of Environment and Conservation of Western Australia (SF008456, SF009067, BB003100, BB003163, BB003312), Department of Parks and Wildlife of Western Australia (08-001825-1) and Parks and Wildlife Commission of the Northern Territory (51604, 58510)

Capture and sampling carried out by Agriculture Victoria Research was done in accordance with permits by the State Government of Victoria Research Permit under Wildlife Act 1975 (FF380519 Permit No: 10004073, FF383165 Permit No: 10005321, FF383294 Permit No: 10006640, FF383493 Permit No: 10007877, FF383578 Permit No: 10008927), and Animal Ethics Research Project Permit (AEC 2019-04). Sampling undertaken by the Northern Australia Quarantine Strategy was undertaken in accordance with a Licence to take Fauna (SF006970) from the Department of Environment and Conservation (WA) and permits from Department of Agriculture, Forestry and Fisheries (QLD – now Department of Agriculture and Fisheries) (CA 2013/07/703) and Department of Agriculture and Fisheries (QLD) (CA 2016/07/980). Permits for collection of faecal samples were not required from Parks and Wildlife (NT) or DAFWA (WA).

For samples collected by Department of Primary Industries, Parks, Water and Environment, Tasmania and Primary Industries and Regions, South Australia, Department of Primary Industries and Regional Development, Western Australia; James Cook University; NSW Department of Primary Industries; University of Technology Sydney; or Biosecurity Queensland, Department of Agriculture & Fisheries, permits were not required for the collection of environmental faecal samples or for samples collected opportunistically from carcasses.

Cloacal samples collected from a wild bird as part of a mortality event investigation by Department of Primary Industries and Regional Development, Western Australia, is also exempt from a permit.

### Sample collection and screening

All samples were collected from wild birds or from wild bird faeces since 2006, as part of the National Avian Influenza Wild Bird Surveillance Program (NAIWB). Details of sample collection and screening methods can be found in [25]. No HPAIV were detected in wild birds through the duration of this study.

### Next generation sequencing

Viral RNA was extracted with MagMAX™-96 viral RNA isolation kit (Thermo Fisher Scientific, Waltham, MA) from avian faecal swab samples, avian swabs and embryonated chicken egg isolated virus samples according to manufacturer’s instructions. Positive samples with an influenza A matrix gene qPCR Ct of ≤30 were selected for influenza A virus targeted next generation sequencing (NGS). The AIV genome segments were amplified using the SuperScript™ III one-step RT-PCR system with high fidelity Platinum™ Taq DNA polymerase (Thermo Fisher Scientific) and universal influenza A virus gene primers as previously described [73]. Sequencing was performed on the Illumina MiSeq NGS platform (Illumina, San Diego, CA) with up to 24 samples pooled per sequencing run by use of dual-index library preparation and the Nextera XT DNA Library Preparation kit and 300-cycle MiSeq Reagent v2 kit (Illumina), according to manufacturer’s instructions. Sequence reads were trimmed for quality and mapped to respective reference sequence for each influenza A virus gene segment using Geneious Prime software (www.geneious.com) (Biomatters, Auckland, NZ).

For a small subset of AIV sequences generated by Agriculture Victoria (Table S1), RNA was extracted using QIAamp Viral RNA Mini Kit (Qiagen, Hilden, Germany), AIV genome amplified [74, 75] and Illumina sequencing libraries prepared using PerkinElmer NEXTFLEX Rapid Directional RNA-Seq Kit 2.0 (Perkin Elmer, Waltham, MA, USA).The libraries were sequenced using a S4 NovaSeq flow cell with the MiSeq 600-cycle v3 kit. The sequences were assembled through the iterative refinement meta-assembler (IRMA) pipeline using the default FLU parameters [76].

### Data availability

Assembled consensus AIV sequences have been deposited in GenBank (accession OL369937-OL372235, OL450375-OL450392) (Table S1).

### Statistical analysis

We analysed sample and sequence metadata for completeness using R 4.0.2 integrated into RStudio 1.3.1073 and the *dpylr()*, *Hmisc()*, *reshape2()*, and *ggplot2()* packages. To compare the differences in Ct values of samples and sequencing “completeness” we used a generalized linear model and a summary of results is presented in Fig S1.

### Phylogenetic analysis

Full-length reference sequences for all AIV segments and subtypes were downloaded from the Influenza Research Database (https://www.fludb.org/). Our sequence search was limited to samples from North America, Europe, Asia and Oceania. Overall, for each of the HA and NA trees, final data sets contained ~500 sequences (+/−20), and for internal segments data sets contained 800-900 sequences. For the internal segment sequences, reference sequences did not include the poultry adapted subtypes H5N1, H7N9, H9N2 or other AIV sequences from poultry. In addition to sequences from Australia generated in this study, we also included all sequences from Oceania (Australia and New Zealand) in GenBank, including partial sequences. Australian H10 sequences [30] that were not available in the Influenza Research Database or GenBank were downloaded from GISAID (https://www.gisaid.org).

Sequences were aligned using MUSCLE v3.8.425 [77] integrated within Geneious Prime. Sequence alignments were cleaned to remove any obviously problematic sequences, including those containing many ambiguous bases, insertions or deletions, and respective data sets were trimmed. Global phylogenetic trees were estimated using the maximum likelihood (ML) method incorporating the most appropriate model of nucleotide substitution estimated using Smart Model Selection in PhyML v3.0 [78, 79]. Trees were visualised using FigTree v1.4.4 (http://tree.bio.ed.ac.uk/software/figtree/). From these global trees we were able to infer the number of independent introductions into Australia, as well as the number of local genome constellations, and used this information to assess the pattern and frequency of segment reassortment.

### Time-scaled phylogenetic analysis

Time-structured phylogenetic trees of all contemporary (those lineages circulating in 2005 or later) Australian lineages of HA, NA and nucleoprotein (NP) sequences, were estimated using the Bayesian Markov chain Monte Carlo method available in BEAST 1.10.4 [80]. Prior to the BEAST analysis ML trees were used to determine the degree of clock-like behaviour of each data set by performing linear regressions of root-to-tip distances against year of sampling using TempEst [81]. All data sets exhibited a strong positive correlation between genetic divergence and sampling time, with correlation coefficients ranging from 0.8-0.99 and R^2^ values ranging from 0.66-0.99. Using BEAST, time-stamped data were analysed under the uncorrelated lognormal relaxed molecular clock [82] and the SRD06 codon-structured nucleotide substitution model [83]. We selected the uncorrelated lognormal relaxed clock following comparisons of the marginal likelihood of the strict and uncorrelated lognormal relaxed molecular clocks for a subset of trees (H4, H5, N6, N8, and two NP lineages) using path/stepping-stone sampling [84]. The Bayesian skyline coalescent tree prior was used as this likely reflects the complex epidemiological dynamics of AIV [85]. Three independent analyses of 100 million generations were performed, which were then combined in LogCombiner v1.8 following the removal of a 10% burn-in. Convergence was assessed using Tracer v1.6 (http://tree.bio.ed.ac.uk/software/tracer/). Maximum credibility lineage trees were generated using TreeAnnotator v1.8 and visualized in FigTree v1.4.

### AIV phylogeography

We selected the HA, NA and NP internal segments as representatives to investigate the phylogeography of AIVs in Australia. Importantly, the selected HA or NA subtypes comprised Australia-specific lineages with >20 sequences (containing sequences from no other continent). For the NP segment, we selected two lineages that comprised only sequences from Australia. Discrete trait analysis was performed using the asymmetric substitution model, and social networks were inferred with Bayesian Stochastic Search Variable Selection (BSSVS) [86]. The extent and pattern of virus movement between locations were determined using Bayes Factor analysis generated by SpreaD3 [87]. We considered Bayes Factors of greater than 10 to be strong support of virus movement between the locations sampled, and greater than 100 to be decisive support [88, 89] within the necessary constraints imposed by sampling bias. The mean number of migration events were inferred by logging/counting the transitions between states along the phylogenetic branches (Markov Jumps) [90]. We also calculated the time spent in the states between two transitions (Markov Rewards) to ensure that rewards were not strongly correlated with export events, thus providing some insight into the effect of sampling bias in our dataset.

## Acknowledgements

The authors would like the contribution of the following individuals: William Steele, Richard Akers, Suelin Haynes and Nick Crosbie from Melbourne Water; Zachary Powell, Heath Dunstan, Lee Manning, John Turnbull, Octavian Manescu, and Mark Jones from Game Management Authority; Jana Batovska, Peter Mee, Xinlong Wang and Macy Hamilton-Smith from Agriculture Victoria Research; Clive Minton, Rosalind Jessop, Robyn Atkinson and Rob Patrick from the Victorian Wader Study Group; Emma Watkins, Marianne Douglas, Teresa Wilson and Kate Swift from Department of Primary Industries, Parks, Water and Environment, Tasmania; Sam Hair, Jamie Ong, Andrew Hughes, Mark O’Dea, Vanessa Rushworth, Emily Glass & Kristine Rayner from Department of Primary Industries and Regional Development, Western Australia; the team from The Northern Australia Quarantine Strategy; K Edla Arzey from NSW Department of Primary Industries; Mick, Todd, Fred van Gessel, Greg Little and Tyler Williams for NSW sampling; Sandy Adsett, Ibrahim Diallo and Allison Crook from Biosecurity Queensland; Kim Critchley from Primary Industries and Regions, South Australia; Chris Bunn and Lyndel Post from the Australian Government Department of Agriculture; Leesa Haynes, Rupert Woods from Wildlife Health Australia (previously Australian Wildlife Health Network); Jemma Bergfeld, Mark Ford, Gemma Harvey, Tristan Reid, Songhua Shan, Som Walker, Jianning Wang, James Watson, David Williams and other diagnostic staff from the CSIRO Australian Centre for Disease Preparedness; Ankita George and Malet Aban from the WHO Collaborating Centre for Reference and Research on Influenza, and staff from the Bolivar Waste Water Treatment Plant. We thank Sebastian Duchene for advice on executing the Markov Jump analysis. In addition to those identified above, the authors would like to thank all individuals, organisations and laboratories that contribute to the NAIWB Surveillance

Program via providing expert input, logistic support and/or collection, analysis and submission of wild bird influenza virus samples and data, including Birdlife Australia, state and territory bird banding groups, wildlife/shorebird groups and hunter groups. We gratefully acknowledge the authors from the originating laboratories and submitting laboratories where genetic sequence data (EPI397842-64) were generated and shared via the GISAID Initiative.

## Supporting information captions

Figure S1. The effect of Ct value of original samples on sequencing success

Figure S2. Temporal distribution of sampling dates.

Figure S3. Phylogeny of H1

Figure S4. Phylogeny of H2

Figure S5. Phylogeny of H3

Figure S6. Phylogeny of H5

Figure S7. Phylogeny of H6

Figure S8. Phylogeny of H7

Figure S9. Phylogeny of H8

Figure S10. Phylogeny of H9

Figure S11. Phylogeny of H10

Figure S12. Phylogeny of H12

Figure S13. Phylogeny of H13

Figure S14. Phylogeny of H16

Figure S15. Phylogeny of N1

Figure S16. Phylogeny of N2

Figure S17. Phylogeny of N3

Figure S18. Phylogeny of N4

Figure S19. Phylogeny of N5

Figure S20. Phylogeny of N6

Figure S21. Phylogeny of N7

Figure S22. Phylogeny of N8

Figure S23. Phylogeny of N9

Figure S24. Maximum likelihood trees for “internal segments”

Figure S25. Data underlying phylogeography assessments of NP

Figure S26. Bayes factor support for migration events

Figure S27. Markov Rewards for each segment presented in Fig 7

Figure S28. Diversity of the “internal” segments for each sampled location

Table S1. Metadata associated with viral genomes generated in this study

Table S2. Details of mixed viruses detected in this study

